# Novel genomic features in entomopathogenic fungus *Beauveria bassiana*: supernumerary chromosomes and putative virulence genes involved in the infection process of soybean pest *Piezodorus guildinii*

**DOI:** 10.1101/2024.06.10.598397

**Authors:** H. Oberti, L. Sessa, C. Oliveira-Rizzo, A. Di Paolo, A. Sanchez-Vallet, M.F. Seidl, E. Abreo

**Author notes:** Corresponding author: Eduardo Abreo, Héctor Oberti.

## Abstract

Biological control methods involving entomopathogenic fungi like *Beauveria bassiana* have shown to be a valuable approach in integrated pest management as an environmentally friendly alternative to control pests and pathogens. Identifying genetic determinants of pathogenicity in *B. bassiana* is instrumental for enhancing its virulence against insects like the resistant soybean pest *Piezodorus guildinii*. This study focused on comparative genomics of different *B. bassiana* strains and gene expression analyses to identify virulence genes in the hypervirulent strain ILB308, especially in response to infection of *P. guildniii* and growth on hydrocarbon HC15, a known virulence enhancer. Strain ILB308 showed the highest number of virulence-related features, such as candidate virulence proteins, effectors, small secreted proteins, and biosynthetic gene clusters. ILB308 also had a high percentage of unique DNA sequences, including six putative supernumerary scaffolds. Gene expression analysis at 4 days post-inoculation revealed upregulation of known virulence factors, including Tudor domain proteins, LysM motif-containing proteins, and subtilisin-like proteases, and novel genes like secreted effectors and heat-labile enterotoxins. Growth on HC15 led to the upregulation of genes associated with oxidoreductase activity related to cuticular alkane degradation and fermentation metabolism/antioxidant responses in the hemolymph. The presence of supernumerary chromosomes and unique virulence genes in ILB308 may contribute to its higher virulence and could be considered as potential targets for enhancing fungal virulence through genetic manipulation.

**Author Summary:** Understanding mechanisms of virulence and virulence enhancement in *Beauveria bassiana* can lay the basis for the development of improved biocontrol agents. Here we used genomic and transcriptomic approaches to study the infection process of ILB308 -an hypervirulent strain- against insect pest *Piezodorus guildinii*. We found that this strain has putative supernumerary chromosomes and an enriched set of presumed virulence proteins like secreted effectors. We infer that the particular assortment of strain specific genes and supernumerary chromosome/s play a role in the degree of virulence exhibited by ILB308 and could be used for future strain improvement strategies.

## Introduction

Insect pests pose a substantial threat to agricultural productivity and consequently to food security and economic stability worldwide [1]. Soybean *Glycine max* (L.) Merril is one of the most economically important crops in the southern United States, Argentina, Brazil, and Uruguay [2], with 371.3 million metric tons projected to be produced in 2030 [3]. The red-banded stink bug *Piezodorus guildinii* (Westwood) (Heteroptera: Pentatomidae) is one of the most detrimental insect pests to soybean in these countries [4,5]. *P. guildinii* direct damage to the crop is caused by direct feeding on the grain during the filling period, which reduces grain yield and seed quality [4,6]. Currently, control methods predominantly rely on the application of insecticides including pyrethroids and neonicotinoids [7,8], which are applied several times during the grain filling period. This practice has led to human and environmental hazards and the development of insecticide resistance, demanding novel approaches for pest management [9,10].

Integrated pest management (IPM) is an environmentally friendly alternative to control pests and pathogens [11]. Biological control methods involving entomopathogenic fungi like *Beauveria bassiana* have shown to be a valuable approach in IPM (Skinner et al., 2014). *B. bassiana* is capable of infecting a wide range of insect hosts [12] including Lepidoptera, Coleoptera, Heteroptera and Thysanoptera [13], including *P. guildinii* [14,15]. Infection can occur via the cuticular route but alternatively it can also enter the insect via the oral route [16,17]. This dual ability to infect insects makes this species a great candidate to develop a biological alternative to chemical pesticides.

Typically, the cuticular infection process of *B. bassiana* can be delimited in different stages covering host recognition, attachment of the spores to the epicuticular surface, penetration of insect cuticle, colonization of the insect hemolymph by hyphal bodies, insect death, and fungal emergence at the exterior for further dispersal (Harith-Fadzilah et al., 2021; Mannino et al., 2019). Efforts have been made to understand and to identify genes involved in the different stages of the fungal infection process [19–22]. Transcriptomic approaches have shown that the infection process is mediated by the co-expression of multiple genes at successive stages of the infection process [23,24]. For instance, it has been proposed that at the early stages of infection genes involved in host cuticle penetration and metabolic processes including amino acid degradation, amino acid/ion transportation, and the secretory system are more relevant while RNA processing, iron transportation, and mitochondrial metabolism processes are more relevant at hyphal bodies inside the host hemolymph [24].

Virulence is a key trait in *B. bassiana* [25,26], which can be enhanced by growing *B. bassiana* on culture media supplemented with insect cuticle components [27–29] or by consecutive inoculations on the target insect [30,31]. However, outcomes vary among tested fungal strains. For example, we have previously examined three *B. bassiana* strains (ILB205, ILB299, and ILB308) and observed differences in virulence towards *P. guildinii*, with ILB308 being the most virulent strain [14]. Pre-culture on n-pentadecane (HC-15) supplemented media, a major volatile compound of *P. guildinii* epicuticle [32], showed an increased virulence in strains ILB299 and ILB308, but not in strain ILB205 [14]. These differences could be related to genetic diversity of specific genes [19], strain-specific features such as supernumerary chromosomes, i.e., chromosomes dispensable for fungal survival but encoding for genes required for specialized functions such as virulence [33], or other strain-specific genes that are not required for growth under all conditions but confer a fitness and adaptive advantage [34,35]. Therefore, understanding the differential responses of strains to different stimuli and elucidating the molecular mechanisms underlying virulence and virulence enhancement is crucial for strain improvement to develop more effective biocontrol agents in the future.

A previous expression analysis of *B. bassiana* strains ILB299 and ILB308 during axenic growth on *P. guildinii* epicuticle, a mimic of the early infection stages, showed a distinct expression pattern for each strain. ILB299 had a higher expression of cuticle penetration-related genes, while ILB308 showed higher expression of cell wall remodelling and cell cycle genes, as well as cuticle adhesion genes [36]. This distinct expression patterns suggest that potentially differences in virulence and HC15 enhancement capacity in *B. bassiana* could be mediated by its genetic background. In this work we performed a genomic comparative analysis of three *B. bassiana* strains. We subsequently focussed on the hypervirulent strain ILB308 to identify core and supernumerary chromosomes, infection process related genes, as well as gene expression changes during the infection of the host insect *P. guildinii*. Furthermore, we sought to identify novel genes involved in virulence of this strain.

## Results

### Assembly of high-quality genomes of three Beauveria bassiana strains

*B. bassiana* strain ILB308 is reported to be more virulent towards *P. guildinii* compared with ILB299 and ILB205, and virulence of ILB308 towards *P. guildinii* can further be enhanced by growth on HC15 [14]. To start determining the genetic underpinnings of these differences in virulence, we sought to compare the genomes of these three strains. First, we obtained a total of 1.3 Gigabases (Gb), 2.9 Gb, and 3.0 Gb of long-read Oxford Nanopore (ONT) data for *B. bassiana* strains ILB205, ILB299, and ILB308, respectively. Based on the genome size of the *B. bassiana* strain ARSEF8028 reference genome assembly of 36Mb [37], we estimated a 36x, 80x and 83x coverage depth for each strain. We complemented these data with 4.1Gb Illumina paired-end (PE) reads for ILB205, 4.2Gb for ILB299, and 4.1Gb for ILB308, providing a mean coverage of 113x per strain. Using an assembly index score based on seven genome assembly statistics, we compared results obtained for four different genome assemblers (SPAdes, Fly, MaSURca, and Canu) and six approaches using Illumina short reads and ONT data. Based on the assembly index score, we determined the best assembly strategy for each strain (**Supplementary File, Table S1-3**), and ultimately selected Flye+POLCA+Samba for ILB205, Masurca for ILB299, and Flye+POLCA for ILB308; for each strain an hybrid assemblies approach with ONT+Illumina data was optimal. For each selected genome assembly, we subsequently removed low-coverage scaffolds and manually incorporated telomere sequences, when possible. All selected methods yielded final genome assemblies with more than 99% complete BUSCO genes. The final genome assemblies were contained in 22 (ILB205 and ILB308) and 33 (ILB299) scaffolds with a total genome size varying between 32.04 to 34.02 Mb (**Supplementary File, Table S1-3)**. Further examination of the genome assemblies revealed a total of 11 telomeric sequences with more than two repetitions of the fungal telomeric repeat sequence ‘TTAAGG’ in ILB205; one scaffold likely represents a complete chromosome with telomeric repeats on both ends. In ILB299 and ILB308, we identified six and three telomeric sequences, respectively, with one scaffold in ILB299 likely representing a complete chromosome. Altogether, we obtained a highly complete and contiguous genome assembly for each *B. bassiana* strain (**Figure 1A**).

**Figure 1.**
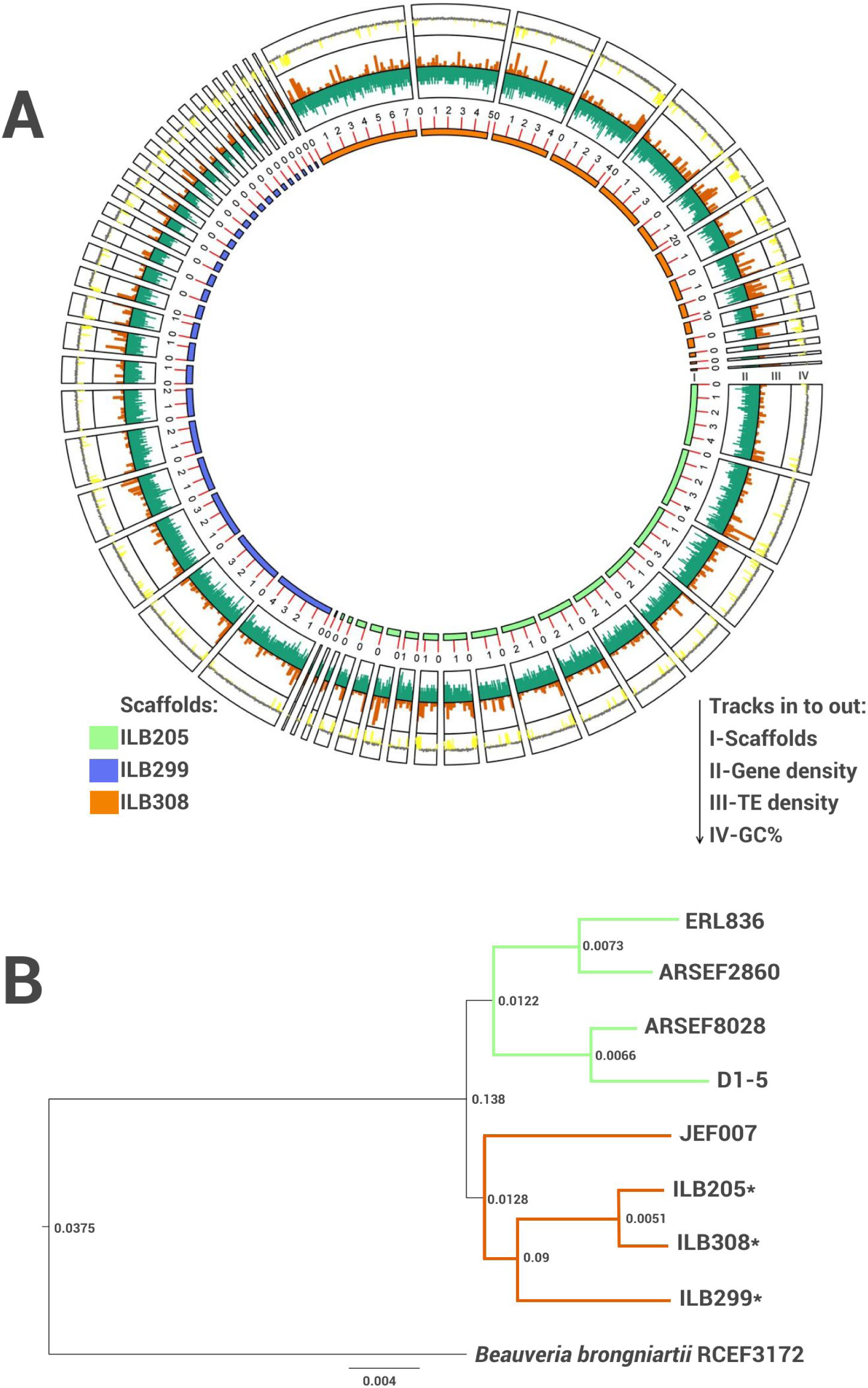
Genome architecture of assembled *B. bassiana* strains and their relationships with publicly available *B. bassiana* strains. A) Circos plot shows key genomics features of *B. bassiana* strains ILB205, ILB299 and ILB308. From inside out: I) Scaffolds longer than 100kb. Red marks in scaffolds represent 1Mb; II) Gene density (genes/10 kb), III) Transposable elements (TE) density (TE/10 kb); IV) GC% (yellow <50% and grey > 50%). B) Phylogenetic tree based on species tree inference from all genes (STAG) using 6,028 single-copy genes of eight *B. bassiana* strains, together with the outgroup *B. brongniartii*, revealed two main clusters: Cluster 1 (green) with four *B. bassiana* strains and cluster 2 (brown) with our three strains (marked with *) and JEF-007. STAG support values are presented at internal nodes and branch lengths represent substitutions per site.

### Structural annotation and phylogenetic relationship between Beauveria bassiana strains

Structural annotation of repeats and protein-coding genes was conducted using homology and *de novo* annotation using the Funannotate pipeline, which yielded a total of 10,763 predicted protein-coding genes for ILB205, 10,555 for ILB299, and 11,330 for ILB308. We then sought to identify core *Beauveria* spp. genes to establish phylogenetic relationships among strains. All protein-coding genes of available *B. bassiana* strains and the outgroup species *Beauveria brongniartii* were used for identifying single-copy orthologous genes using Orthofinder2. A total of 7,549 core *Beauveria* spp. genes were detected, 6,028 of which were single copy. The resulting dataset was used for the construction of a maximum-likelihood phylogenetic tree, which uncovered two main clusters of *B. bassiana* strains (**Figure 1B**). One of these clusters is formed by four strains isolated and reported from different geographical locations including strains ARSEF8028 (Denmark), ARSEF2860 (USA), D1-5 (China), and ERL826 (South Korea). The other cluster was formed by the three *B. bassiana* strains analyzed in this study (all from Uruguay) and the strain JEF-007 (South Korea) (**Figure 1B**).

To further explore the genomic diversity of the *B. bassiana* strains, we performed pairwise genome-wide comparisons between ILB205, ILB299, ILB308, and the reference genome assembly ARSEF8028 (**Table 1**). As expected, given the phylogenetic relatedness between the three *B. bassiana* strains from Uruguay, we observed a high degree of genomic coverage (65.0 - 93.4%), based on ≥75% sequence identity, but not with strain ARSEF8028 (16.1-26.7%). Moreover, in line with the phylogenetic analysis, ILB205 and ILB308 share the highest coverage (93.4%) (**Table 1**; **Figure 1B**).

**Table 1.** Percentage of genome coverage with similarity between *B. bassiana* strains.

|  |  | Strain |  |  |  |
| --- | --- | --- | --- | --- | --- |
|  | Query | ILB205 | ILB299 | ILB308 | ARSEF8028 |
| **>75% similarity** | ILB205 | 100 |  |  |  |
|  | ILB299 | 85,9 | 100 |  |  |
|  | ILB308 | 93,4 | 65,0 | 100 |  |
|  | ARSEF8028 | 16,1 | 26,7 | 18,2 | 100 |
| **0% similarity** | ILB205 | 0 |  |  |  |
|  | ILB299 | 2,8 | 0 |  |  |
|  | ILB308 | 2,1 | 1,5 | 0 |  |
|  | ARSEF8028 | 7,5 | 6,6 | 10,3 | 0 |

### Functional predictions reveal candidate virulence genes in *Beauveria bassiana*

To predict and prioritize genes with potential involvement in virulence in the analyzed *B. bassiana* strains as well as the reference strain ARSEF8028, we functionally annotated the genes using a comprehensive sequence similarity search against the NCBI non-redundant and the eggNOG databases. We functionally annotated a significant proportion of the predicted proteomes in ILB205, ILB299, ILB308, and ARSEF8028, amounting to 68.8%, 69.6%, 67.2%, and 71.42% of their total protein-coding genes, respectively (**Table 2**). We also predicted the presence of biosynthetic gene clusters (BGC) encoded in each *B. bassiana* strain **(Supplementary File Table 4)**, with ARSEF2860 encoding the highest number followed by ILB308 (**Table 2**). Importantly, the four analyzed strains encode three well known *B. bassiana* virulence BGC, oosporein, beauvericin, and bassianolide (Ortiz-Urquiza and Keyhani, 2013).

**Table 2.**
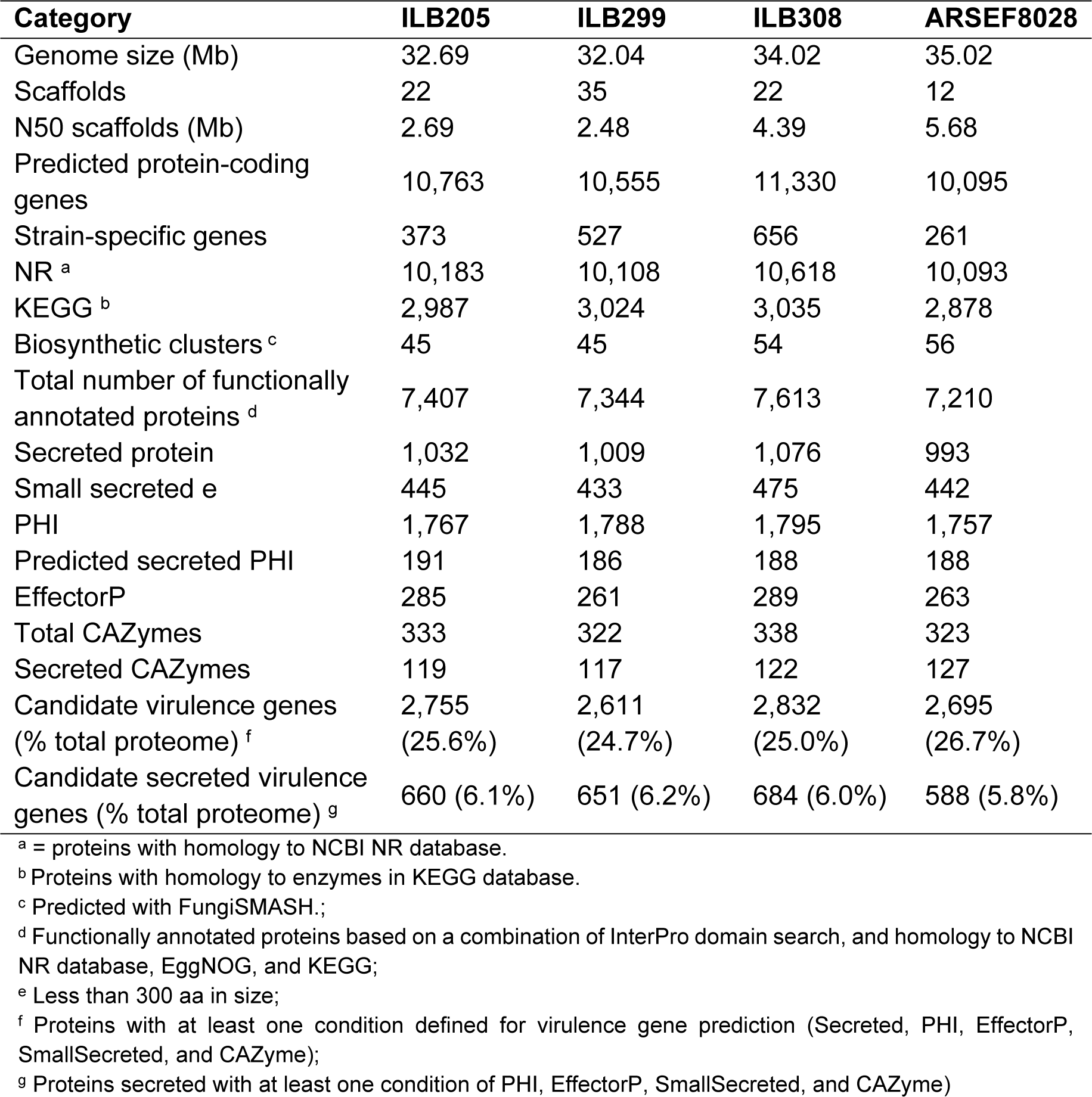
Summary of structural and functional annotation and overview of candidate virulence genes identification in *Beauveria bassiana* strains.

| Category | ILB205 | ILB299 | ILB308 | ARSEF8028 |
| --- | --- | --- | --- | --- |
| Genome size (Mb) | 32.69 | 32.04 | 34.02 | 35.02 |
| Scaffolds | 22 | 35 | 22 | 12 |
| N50 scaffolds (Mb) | 2.69 | 2.48 | 4.39 | 5.68 |
| Predicted protein-coding genes | 10,763 | 10,555 | 11,330 | 10,095 |
| Strain-specific genes | 373 | 527 | 656 | 261 |
| NR <sup>a</sup> | 10,183 | 10,108 | 10,618 | 10,093 |
| KEGG <sup>b</sup> | 2,987 | 3,024 | 3,035 | 2,878 |
| Biosynthetic clusters <sup>c</sup> | 45 | 45 | 54 | 56 |
| Total number of functionally annotated proteins <sup>d</sup> | 7,407 | 7,344 | 7,613 | 7,210 |
| Secreted protein | 1,032 | 1,009 | 1,076 | 993 |
| Small secreted <sup>e</sup> | 445 | 433 | 475 | 442 |
| PHI | 1,767 | 1,788 | 1,795 | 1,757 |
| Predicted secreted PHI | 191 | 186 | 188 | 188 |
| EffectorP | 285 | 261 | 289 | 263 |
| Total CAZymes | 333 | 322 | 338 | 323 |
| Secreted CAZymes | 119 | 117 | 122 | 127 |
| Candidate virulence genes<br>(% total proteome) <sup>f</sup> | 2,755<br>(25.6%) | 2,611<br>(24.7%) | 2,832<br>(25.0%) | 2,695<br>(26.7%) |
| Candidate secreted virulence<br>genes (% total proteome) <sup>g</sup> | 660 (6.1%) | 651 (6.2%) | 684 (6.0%) | 588 (5.8%) |

<sup>a</sup> = proteins with homology to NCBI NR database.
<sup>b</sup> Proteins with homology to enzymes in KEGG database.
<sup>c</sup> Predicted with FungiSMASH;
<sup>d</sup> Functionally annotated proteins based on a combination of InterPro domain search, and homology to NCBI NR database, EggNOG, and KEGG;
<sup>e</sup> Less than 300 aa in size;
<sup>f</sup> Proteins with at least one condition defined for virulence gene prediction (Secreted, PHI, EffectorP, SmallSecreted, and CAZyme);
<sup>g</sup> Proteins secreted with at least one condition of PHI, EffectorP, SmallSecreted, and CAZyme)

We then identified candidate virulence proteins (CVP) and secreted CVP (SCVP) using five *in silico* approaches based on homology searches and *ab initio* prediction software. Interestingly, the four *B. bassiana* strains have a similar number and proportion of these genes (**Table 2**). Although ILB308 does not show a higher proportion of CVP compared to the other three strains, it contains a higher number of genes in this category (**Table 2**), which might contribute to its higher virulence.

### Hypervirulent strain ILB308 harbors strain specific genes and putative supernumerary chromosomes that potentially contribute to higher virulence

*B. bassiana* has been proposed to have strain specific genomic regions like supernumerary chromosomes (Eivazian-Kary and Alizadeh, 2017), genomic features that could be involved in fitness and niche adaptation in entomopathogens [39]. Thus, it is conceivable that part the observed virulence differences of ILB308 with ILB205 and ILB299 strains could be related to these genomic features. Notably, 2.1%, 1.5%, and 10.3% of ILB308 genomic sequences were absent in ILB299, ILB205, and ARSEF8028, respectively (**Table 1**). Moreover, based on gene families identified by Orthofinder2, 656 strain-specific genes (SSG) were identified in ILB308 and 44 of these genes were also annotated as CVP. A significant proportion (39%) of these 44 genes do not show any homology to proteins of other organism in the non-redundant sequence database, suggesting that these genes could be associated to specific ILB308 traits like virulence in *P. guildinii*.

To identify potential supernumerary chromosome(s) in our genome assemblies, we combined multiple *in silico* criteria like presence of species single-copy core genes, GC%, TE content, and sequencing read coverage. We identified a set of 13 scaffolds (scaffolds 1 to 12 and 17) that most likely represent core genomic regions as these harbors in total 6,351 single-copy *B. bassiana* core genes (**Supplementary File, Table 5)**. By contrast, scaffolds 22 and 25 did not encode any protein-coding genes, and we thus excluded these from further analyses. Principle component analysis (PCA) and hierarchical clustering were used to classify all scaffolds, and we identified non-core, single-copy gene scaffolds (scaffolds 13-16, 19, and 20) that exhibited distinct genomic patterns compared with the putative core scaffolds (**Figure 2A**). To investigate the presence of these specific scaffolds in the other two strains (ILB205 and ILB299), we determined the sequencing read coverage over the core and putative supernumerary scaffolds based on mapping of the ONT data derived from the three *B. bassiana* strains (**Figure 2B**). As expected, core scaffolds are contiguously covered by sequencing reads from all three strains (**Supplementary Figure 1**), while scaffolds 13-16, 19, and 20 show multiple regions that lack sequencing coverage (**Figure 2B**). Most regions with lack of coverage corresponds to regions with transposable elements or unique genes, while regions that do show mapping with ILB205 and ILB299 data possibly correspond to duplicated genes from core scaffolds that are present in these two strains. Based on these analyses, we conclude that these six scaffolds likely represent regions of (a) putative supernumerary chromosome(s) in ILB308, which might have implication on the distinct virulence profile of ILB308. Putative supernumerary scaffolds range in size from 11 to 143 kb, and contained a total of 157 genes, of which most of these genes (100) where duplications of genes present in core scaffolds (**Supplementary File, Table 6)**. Moreover, a total of 46 (29%) were annotated as SSG and 19 of these 46 were present exclusively in these supernumerary scaffolds and thus absent in the core (**Supplementary File, Table 6)**. Interestingly, there is a significant enrichment of SSG in supernumerary scaffolds compared with core scaffold (29 % vs 5.4%), which reinforces the idea of the presence of genes related to specific adaptations residing in the supernumerary chromosomes of ILB308.

**Figure 2.**
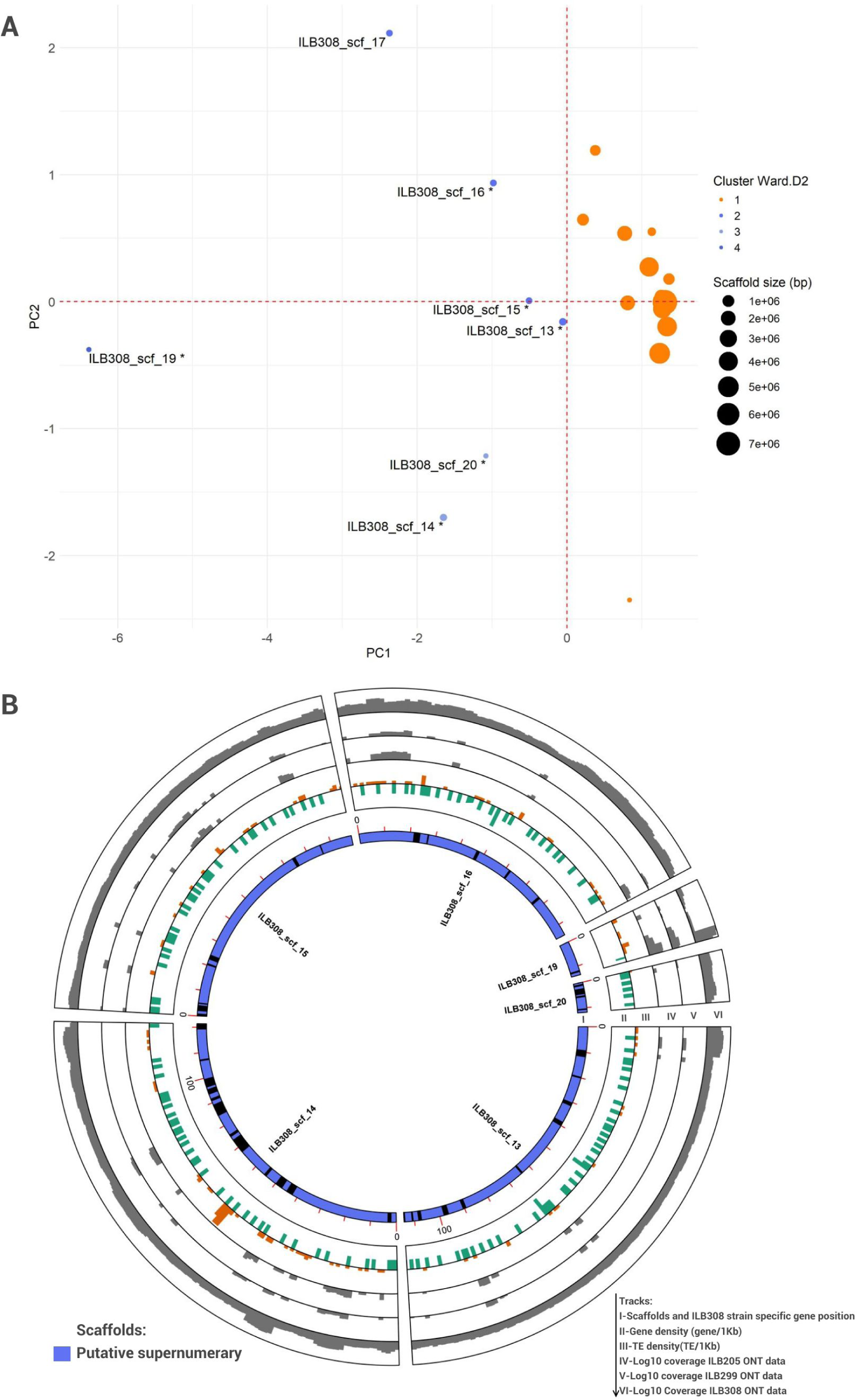
*B. bassiana* strain ILB308 harbors supernumerary scaffolds with high proportion of strain-specific genes. A-Principal component analysis (PCA) shows a distinct separation between core and putative supernumerary scaffolds in ILB308. Cluster 1 shows putative core scaffolds and Cluster 2-4 putative supernumerary scaffolds. Labels only added for Clusters 2-4. * Indicates scaffolds that lack core single-copy *Beauveria spp.* genes. B-Circos plot of the six putative supernumerary scaffolds (blue). Tracks I-Scaffolds and ILB308 strain-specific gene position, II-Gene density (number of genes/1kb); III-TE density (number of TE/1kb); IV-SSG position; V-Coverage of strain ILB205 Oxford Nanopore (ONT) data (Log10); VI-ILB299 ONT data Log10 coverage; VII-ILB308 ONT data Log10 coverage.

### GFP-labelled ILB308 strain and infection progression in *Piezodorus guildinii*

To investigate the infection process of hypervirulent strain ILB308 in *P. guildinii*, we constructed a GFP-labeled transformant. The binary vector pCGEN-GFP was used for *Agrobacterium tumefaciens* mediated transformation. We evaluated the radial growth (**Supplementary Figure 2A**) and the virulence (**Figure 3A**) of wild type and GFP-labelled strains and verified the GFP signal in the transformant (**Supplementary Figure 2B**). Based on this we demonstrated that these two strains do not show any phenotypic and virulence differences. The infection process in *P. guildinii* adults was monitored daily. No deaths were observed in adult insects between 1-3 dpi (**Figure 3A**), which coincided with a low GFP signal (not significant) throughout the body regions of *P. guildinii* at these time points (**Figure 3B**). Based on these results, we suggest that early stages of infection (host recognition, adhesion, and cuticle penetration) occur between 1-3 dpi. At 4 dpi, we detected an intense GFP signal, and survival rate of the insect was reduced to 90%. Based on this, we concluded that successful penetration had occurred, and that the colonization of the hemolymph was beginning at 4 dpi. We consequently selected this time point for RNA extraction and subsequent RNAseq analyses.

**Figure 3.**
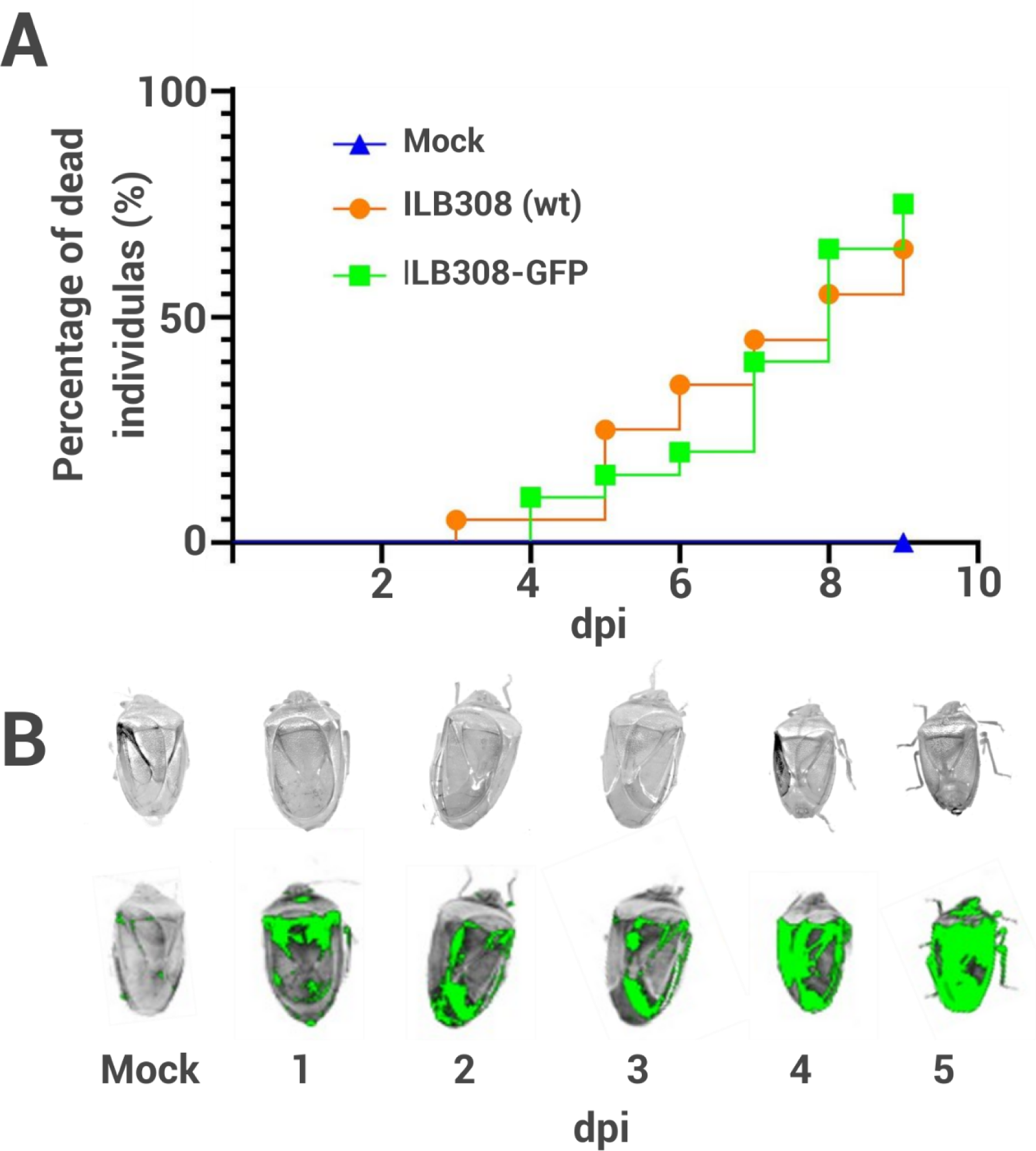
Mortality and fluorescence analyses show that 4 days post inoculation (dpi) is a key timepoint in the infection of *P. guildinii* adults with *B. bassiana* strain ILB308. A) Survival graph of *P. guildinii* insects infected with mock and an GFP-labeled *B. bassiana* strain (ILB308). Line graphs represent the survival percentage at different time points of *P. guildinii* adults treated topically with GFP-labeled ILB308 and mock-treated controls. B) Growth of GFP-labeled ILB308 based on GFP accumulation on *P. guildinii* adults after inoculation using a fluorescent transilluminator at 1 to 5 days post inoculation (dpi). Upper row shows the body outline of *P. guildinii* adults and lower row shows the saturated GFP signal at the same treated insects.

### ILB308 shows expression of putative virulence related proteins and genes present in biosynthetic clusters during infection of *Piezodorus guildinii*

To identify candidate genes that are activated during insect colonization of ILB308 strain, we examined the transcriptome of ILB308 strain at 4 days post-inoculation (dpi) of *P. guildinii*. A total of 30Gb and 42.2Gb of axenic (*B. bassiana*) and infection dual-RNAseq (*P. guildinii* and *B. bassiana* at 4dpi) data was obtained, respectively, and RNA sequencing (RNAseq) reads were mapped to the ILB308 genome assembly (**Supplementary file Table S7)**. For differential expression analysis, pairwise comparisons were made between ILB308 at 4 dpi (infection) with ILB308 grown in minimal media (MM) (axenic). Altogether, we observed a total of 3.796 differentially expressed genes (DEG), of which 1,169 were upregulated during *P. guildinii* infection at 4dpi (**Figure 4A**), representing 10.3% of the total ILB308 predicted transcriptome.

**Figure 4.**
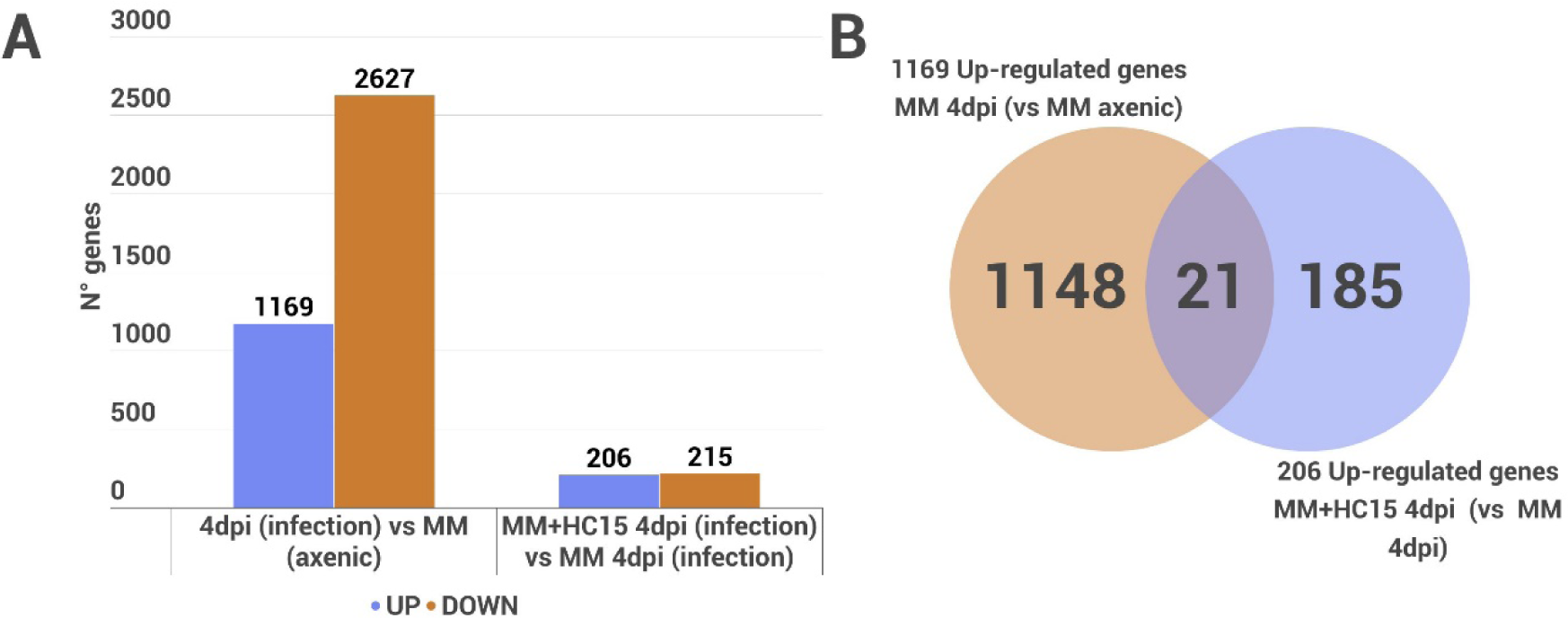
Gene expression analysis shows several differentially expressed genes related to infection process and virulence enhancement in *B. bassiana* strain ILB308 at 4 days post inoculation (dpi). A) Number of differentially expressed genes in *B. bassiana* strain ILB308. Differentially expressed genes were identified at a false discovery rate (FDR) < 0.05 in pairwise comparisons with axenic culture and during infection of *P. guildinii*. B) Number of unique and shared upregulated genes in ILB308 during infection with (MM+HC15 4dpi) and the control (MM without HC15; 4dpi).

The 1,169 upregulated genes during infection were enriched in three KEGG pathways and 378 Gene Ontology (GO) terms (**Supplementary File, Table 8-9**). Two of the three enriched KEGG pathways are “Ribosome” (ko03010) and “Aminoacyl-tRNA biosynthesis” (ko00970), related to genetic information processing, suggesting significant differences in protein synthesis and translation during infection. The enrichment of GO terms such as “cytoplasmic ribonucleoprotein granule”, “cytoplasmic translation,” “peptide metabolic process,” and “translation factor activity, RNA binding” further emphasizes the importance of the cellular translation machinery during the infection process. Additionally, GO terms like “cell cycle DNA replication” suggest active DNA replication. Overall, these enriched pathways and GO terms highlight the dynamic interplay of genetic information processing, translation, and protein synthesis in the strain ILB308 during infection at 4 dpi.

To determine whether known *B. bassiana* virulence genes are upregulated during the interaction with *P. guildinii* at 4 dpi, we searched within the set of upregulated genes for homologs of known virulence factors. We only found homologs to four (*Blys2, Blys5, BbTdp1* and *Pr1C*) out of the 64 known virulence genes (**Supplementary File, Table 10**), suggesting that either most known virulence genes are expressed at a different timepoint in this strain and/or that a different set of virulence related genes, which would be specific to ILB308, are expressed and, subsequently, required for virulence.

Compounds produced by BGC have also been reported to be involved in virulence in *B. bassiana,* including beauvericin [40], bassianolide [41], and oosporein [42]. Five genes of the oosporein cluster (*BBILB308_005136, BBILB308_005130, BBILB308_005141, BBILB308_005135, BBILB308_005131*), two of the beauvericin (*BBILB308_002462, BBILB308_002463*), and one of bassianolide (*BBILB308_007677*) were upregulated during colonization of *P. guildinii* at 4dpi, indicating that a combination of these compounds may potentially play a role in virulence of ILB308. Moreover, genes involved in other BGCs such as beauveriolide (*BBILB308_005089*), choline (*BBILB308_002810, BBILB308_002806*) and metacholine (*BBILB308_008932, BBILB308_008937, BBILB308_008927, BBILB308_008928, BBILB308_008929, BBILB308_008930*) were also upregulated. These results indicate that previously identified *B. bassiana* BGCs are involved in infection of different insects, and our data suggests that these and also additional BGCs not yet studied in detail, might be involved in virulence against *P. guildinii*.

Genes with highest fold changes (FC) in gene expression might play prominent roles during the infection process. Thus, we identified the top-ten DEG with the highest FC at 4dpi compared with MM (**Table 3**). Of these ten genes, none was homologous to known *B. bassiana* virulence genes or BGCs. Interestingly, four out of ten were predicted as SCVP and one of these was also a SSG (**Table 3**). Between these four SCVP genes, *BBILB308_010764* was predicted as a hypothetical, small, secreted effector protein. Moreover, within the top 10, we observed a secreted protease and a Heat-labile enterotoxin, both proteins with known involvement in the infection process of entomopathogenic fungi [17]. Thus, this set of ten genes, and specifically those predicted as SCVP, could represent important virulence factors with yet unknown function.

**Table 3.** ILB308 top ten upregulated genes during infection of *P. guildinii* at 4 dpi.

| ID | NCBI NR description | \|FC\| | *In silico* predictions | Attribute |
| --- | --- | --- | --- | --- |
| *BBILB308_010764* | hypothetical protein BB8028_0003g01300 [Beauveria bassiana ARSEF8028] | 107.2 | Secreted+EffectorP+Small secreted protein. | SCVP |
| *BBILB308_009995* | hypothetical protein BBAD15_g8370 [Beauveria bassiana D1-5] | 91.9 | Secreted | SCVP |
| *BBILB308_009715* | SCF E3 ubiquitin ligase complex F-box protein grrA [Beauveria bassiana ARSEF 2860] * | 69.0 | - | - |
| *BBILB308_000024* | Minor extracellular protease vpr [Beauveria bassiana ERL836] | 29.6 | Secreted | SCVP |
| *BBILB308_004334* | SCF E3 ubiquitin ligase complex F-box protein grrA, BBA_09932 [Beauveria bassiana ARSEF 2860] | 24.9 | - | - |
| *BBILB308_007302* | hypothetical protein BBAD15_g11496 [Beauveria bassiana D1-5] | 24.3 | - | - |
| *BBILB308_002952* | Heat-labile enterotoxin IIA, A chain [Beauveria brongniartii RCEF 3172] | 21.8 | Secreted | SSG, SCVP |
| *BBILB308_010579* | hypothetical protein CRV24_005104 [Beauveria bassiana ERL836] | 19.4 | - | - |
| *BBILB308_007305* | C2H2 finger domain-containing protein [Beauveria bassiana ARSEF 2860] | 17.4 | - | - |
| *BBILB308_011433* | hypothetical protein BB8028_0010g00288 [Beauveria bassiana ARSEF8028] | 12.8 | - | - |

SCVP=Secreted candidate virulence protein; SSG=Strain-specific genes.

### 1.1 Putative supernumerary scaffolds of ILB308 are involved in the infection process of *Piezodorus guildinii*

Based on our gene expression analyses, we showed that twelve genes upregulated at 4dpi (**Supplementary File, Table 11**) were present on three putative supernumerary scaffolds. Five of these genes were single copy SSG (*BBILB308_011322*, *BBILB308_011337, BBILB308_011255, BBILB308_011216* and *BBILB308_011215)* and four CVP (*BBILB308_011217, BBILB308_011218, BBILB308_011220* and *BBILB308_011244).* All five SSG and two out of the four CVP showed homology to hypothetical proteins of others entomopathogenic fungi including *Metarhizium robertsii*, *Cordyceps javanica*, and *Aaosphaeria arxii* (**Supplementary File, Table 11**). Expression of protein-coding genes located in the putative supernumerary scaffolds at 4 dpi suggest that this strain-specific genomic regions could contribute to the pathogenicity toolbox, and especially those genes that are single-copy SSG could be relevant for the specific mechanism of the ILB308 infection process.

### ILB308 growth in n-pentadecane induces expression of genes related to oxidoreductase activity

To determine the expression changes caused by growth of ILB308 on HC15 supplemented medium on gene expression during insect infection, we compared 4 dpi insect samples with ILB308 spores grown in MM or MM supplemented with HC15 (MM+HC15). A total of 206 genes were differentially expressed, of which 79 were classified as CVP and five as SSG (**Supplementary File, Table 12).** Among the five upregulated SSG, one of these genes was also a SCVP, homologue to a Heat-labile enterotoxin IIA, A chain of *Metarhizium brunneum* (*BBILB308_00*2969). One SSG was encoded in one supernumerary scaffold, homologue to a hypothetical protein of *B. brongniartii* (*BBILB308_11285*).

Of the 206 upregulated genes, 21 have been also detected as upregulated during infection (**Figure 4B; Supplementary File, Table 12**). These genes therefore could represent genes important for infection and involved in the enhancement of virulence. Among these 21 shared genes, only one (*BBILB308_005455*) is a known *B. bassiana* virulence factor (*Blys2*) (**Supplementary file, Table 10**). Moreover, “oxidoreductase activity” was the only enriched GO term in these 206 genes, and no enriched KEGG pathways were identified. Among “oxidoreductase activity” we found a total of 35 genes (**Supplementary file, Table 13**), including five Cytochrome P450 monooxygenases, three alcohol dehydrogenase, two Glycerol 2-dehydrogenase, and one aldo-keto reductase. The upregulation of these genes could be related to active alkane degradation, fermentation metabolism, and antioxidant/detoxification, which collectively could contribute to an enhanced virulence in ILB308.

## Discussion

Identification of genetic determinants of pathogenicity in entomopathogenic fungi such as *Beauveria bassiana* is relevant since manipulation of these genes could be a mean to further improve virulence towards insects. This virulence enhancement could be accomplished either by physiological modification of gene expression [14], by genetic modifications like gene overexpression [43–46], or by protoplast fusion [47,48]. Strains with enhanced virulence could contribute to biological control strategy against resistant insects like *P. guildinii*. Thus, we mined for virulence genes in the hypervirulent *B. bassiana* strain ILB308 and in two other strains (ILB205 and ILB299) by comparative genomics and by gene expression analyses of ILB308 in response to infection and infection after growth on HC15, a known virulence enhancer [14].

Structurally, we found that the most virulent strain to *P. guildinii*, ILB308, had the highest numbers of CVP, effectors (EffectorP), small secreted proteins, and BGC. Moreover, we demonstrated that ILB308 has the highest percentage of unique sequences, including the presence of six putative supernumerary scaffolds that could represent parts of one or more supernumerary chromosomes. The presence of supernumerary chromosomes has been previously suggested in *B. bassiana* (Eivazian-Kary and Alizadeh, 2017), but was never addressed with genomic approaches. As reported in other pathogenic fungi including *Fusarium oxysporum* [49,50] and *Nectria haematococca* [33], a high proportion of genes within these supernumerary scaffolds were strain-specific in ILB308 (29%). Moreover, like *Metarhizium* spp. much of these SSG only share homology to proteins of unknown function [51] and are also present in other entomopathogenic fungal species like *A. arxii*, *C. javanica*, or *M. robertsii*. How these supernumerary scaffolds, harboring genes different to other *B. bassiana* strains, were gained or transferred is yet an open question. It is possible that these are transferred between different strains and or species via horizontal transfer, as was reported recently in *M. robertsii* [39]. Different strains that interact during fungal co-infections on the insect cuticle could facilitate the passage of supernumerary chromosomes and may provide fitness benefits in entomopathogens [39]. These results suggest that structural adaptations at both gene and chromosomal levels in ILB308 strain could support its higher virulence. This genomic feature might therefore be valuable for improving biocontrol agents, for instance through the transference of full supernumerary chromosomes to combine different strains to gain or improve virulence.

Beyond the structural features of ILB308, we sought to determine which genes were upregulated during infection and how their expression was altered by growth in the presence of HC15. We selected 4dpi to study gene expression at the infection process through RNA-seq, when an increase of insect mortality and GFP-signal over the insect body coincided. Our expression analysis at 4dpi showed that a few known virulence factors and three well-known BGC *B. bassiana* were upregulated. Specifically, among the upregulated genes during infection, we found homologues genes to *B. bassiana* ARSEF2860 such as the Tudor domain containing protein (*Tdp1*), a gene related to the conidiation process and blastospore proliferation in the hemocoel that is crucial for infection of *G. mellonella* [52]. This gene was also reported as upregulated in ILB308 strain during growth in media supplemented with exuviae of *P. guildinii* in Sessa et al (2024)[36], suggesting that this gene is highly expressed in the first stages of infection and hemocoel colonization by ILB308 in *P. guildinii* and could be a determinant gene for virulence towards this insect. Moreover, we found two homologues to the LysM motif containing proteins *Blys2* and *Blys5* upregulated during infection of ILB308. Both genes code for a secreted effector protein containing LysM domains, a type of protein that has been extensively characterized in plant pathogens as an effector involved in sequestering the chitin oligosaccharides released from the fungal cell walls [53,54]. These two genes were reported to be necessary for the full virulence of *B. bassiana* against *G. mellonella* as they are involved in coating and protecting the cell walls of *B. bassiana* from host cell recognition, delaying the insect defense response [55]. Therefore, co-expression of both genes could also be important during *P. guildinii* colonization, and therefore hint towards an important defense evasion mechanism of *B. bassiana*. Upregulated gene expression of the homologue of the Subtilisin-like *Pr1C* secreted protease was an interesting finding since this gene has been reported in *B. bassiana* as involved in the epicuticle degradation in the early stages of infection in *G. mellonella* larvae [56]. Upregulation of this gene could be related to the active colonization of the external layer of adults *P. guildinii* cuticle as was shown by the increase of GFP signal at 4dpi, and not directly involved in the hemolymph colonization by ILB308. Genes present in BGC of *B. bassiana* involved in infection like oosporein, beauvericin, and bassianolide were also found to be upregulated. These toxins have been reported to help *B. bassiana* to parasitize and kill its hosts [40–42,57]. The presence of these three BGC in ILB308 as well as in ILB299 and ILB205 shows the production of these compounds is most likely common to these strains. Upregulation in ILB308 at 4dpi could be expected since expression of some genes of these BGC during hyphal bodies colonization of hemolymph of *G. mellonella* has similarly been reported [24].

The low number of previously known virulence genes of *B. bassiana* upregulated during infection at 4 dpi suggests that ILB308 strain might have specific mechanisms to invade and colonize *P. guildinii*. To determine novel virulence genes mediating *P. guildinii* infection, we focused on the ten genes with the highest upregulation at 4dpi. We found that two of the top ten expressed genes were SCVP genes with unknown function, and one of them was a predicted small secreted effector protein. Effector proteins are commonly secreted and annotated as hypothetical proteins with unknown function due lack of sequence homology [58]. These proteins are involved in host response evasion and target specific host proteins in plant pathogens [59], but their role in entomopathogen fungi is still unclear. Small secreted proteins are frequently associated with host adaptation or specialization [60], but further functional analysis should focus on these candidate proteins to corroborate their functions during *B. bassiana* infection towards *P. guildinii*. Among these ten top genes, we also found a homologue of a “Heat-labile enterotoxin IIA, A chain”. Heat-labile enterotoxins composed of a subunit A (LTA) are prevalent in the genomes of fungal entomopathogens, and have been recently reported in *B. bassiana* associated with virulence against *G. mellonella* [61]. The gene found in our study shows homology to an LTA from *M. brunneum*. LTA proteins contribute to the fungal-host interaction in *B. bassiana* via maintaining the homeostasis of carbohydrate profiles on the cell surface [61], and these genes particularly could be important during oral toxicity/infection [17]. This type of protein has also been reported as upregulated during blastospore formation in *Metarhizium rylei* [62], a process that occurs within the insect hemolymph. Interestingly, an SSG gene homologue to a *M. brunneum* LTA was also upregulated during infection after growth in HC15, suggesting that this type of proteins can be involved in virulence as well as in increased virulence against *P. guildinii,* yet further analysis needs to be performed to imply these specific proteins to the infection process.

Growth in HC15 supplemented media increases virulence of ILB308 strain as reported previously by Sessa et al., (2022)[14]. We here reported the upregulation of 206 genes during insect infection upon HC15 enhancement. Among these, we noted an enrichment of genes associated with the GO term “Oxidoreductase activity”, which is not unexpected since hydrocarbon assimilation like HC15 produces oxidative stresses with high levels of reactive oxygen species and antioxidant response in fungal cells [63]. These upregulated genes could be roughly subdivided by their main involvement in alkane degradation, fermentation metabolism, or antioxidant/detoxification activity. Regarding alkane degradation, five Cytochrome P450 monooxygenases (CYP450) and one Peroxisomal acyl-coenzyme A oxidase were upregulated. Besides CYP450, which are proteins known to be involved in alkane degradation [28] and enhanced after growth on HC15 [36], it is noteworthy that an acyl-coenzyme A oxidase was found to be upregulated. This enzyme is active in the n-alkane-assimilating yeast *Yarrowia lipolytica* [64] and a significant increment in acyl-CoA oxidase activity can be observed in peroxisome fractions of *B. bassiana* grown either on n-C16, n-C24 or n-C28 alkanes as compared with the same fraction of glucose-grown fungi [63]. Therefore, at 4 dpi, pre-growth on HC15 could be associated with enhanced hydrocarbon assimilation and improved resource acquisition by the fungus. Interestingly, we also found at 4dpi that genes involved in fermentation of sugars (alcohol dehydrogenases), glycerol (glycerol dehydrogenase), and lactate (FMN-dependant dehydrogenase functionally annotated as lactate dehydrogenase) were upregulated. In this regard, *B. bassiana* growth in the haemolymph of ticks was reported to prompt changes like sugar depletion, glycerol mobilization from the tick tissue, and lactic acid accumulation, which result from switching from aerobic to anaerobic or microaerobic conditions [65,66]. This scenario is consistent with the observed upregulation of fungal alcohol, glycerol, and lactate dehydrogenase in ILBB308 at 4 dpi found, when the fungus needs to obtain energy from alternative pathways and variable sources in the haemolymph. In addition, the importance of alcohol dehydrogenase in fungal infection has been demonstrated in entomopathogen *Metarhizium anisopliae* through the knockdown of alcohol dehydrogenase *ADH1* and resultant decrease in virulence [67]. Besides the nutritional shifts that condition the metabolic pathways of the fungus, the insect also produces toxic compounds like reactive oxygen species to counteract fungal infection [68,69]. Therefore, the observed upregulation of oxidoreductase like aldo-keto reductase following growth on HC15, an oxidative stressor, and during infection is plausible. Aldo-keto reductases in fungi are known for their involvement in metabolic adaptation, secondary metabolism [70], oxidative response [71], and are associated with detoxification processes [72]. For example, in *B. bassiana*, an aldo-keto reductase, *Bbakr1,* was identified as being involved in stress response and the detoxification of heavy metal chromium, although it was not required for virulence in *B. bassiana* [73]. It is therefore possible that, whereas not being essential for virulence, its upregulation still exerts a positive effect on fungal infection in the stressful conditions of the haemolymph. In summary, growth in HC15 seems to pre-activate a diverse set of genes which can have a positive effect during the advanced stages of the infection process as well as during epicuticle penetration [14]. How these genes are activated by growth in HC15 is unclear and needs to be addressed in the future.

In conclusion, we here performed comparative genomic and expression analyses and uncovered genetic determinants of the virulence of *B. bassiana* strain ILB308 towards *P. guildinii*. This information provides the necessary framework to aid the development of improved fungal strains that can be exploited as biopesticides. Importantly, we here computationally identified supernumerary chromosomes and novel candidate genes that are likely involved in the *B. bassiana* infection process. Functional verification of their roles during infection and their specific functions will need to be addressed in future works.

## Materials and Methods

### Biological materials

*Beauveria bassiana* strains ILB205, ILB299, and ILB308 were obtained from the INIA Las Brujas fungal collection (WDCM 1291), Uruguay. The strains were grown on potato dextrose agar (PDA) plates at 26°C. These three strains exhibit varying degrees of virulence towards *P. guildinii* [14], with ILB308 strain being the most virulent, followed by ILB205 strain with medium virulence, and ILB299 with the lowest virulence.

*Piezodorus guildinii* individuals were obtained from an indoor mass rearing and were maintained at 24 ± 1 °C, 80 ± 10% RH, 14/10 h of light/dark on fresh soybeans, green beans, and water for two generations. Adult individuals of 2-3 weeks were used for all inoculation assays.

### DNA purification and whole-genome sequencing

Vegetative mycelia were harvested from cultures of strains ILB205, ILB299, and ILB308 grown on PDA for 1 week at 26°C. Fungal genomic DNA was extracted using the GeneElute kit (Qiagen, Germany) following the manufacturer guidelines. The DNA samples were utilized for genome sequencing using two distinct sequencing platforms. Illumina short reads of 150 base pairs (bp) in paired-end (PE) format were sequenced using the Novaseq 6000 platform (Illumina, San Diego, CA) at Novaseq Inc, USA. Long reads were generated using SQK-LSK112 Ligation Sequencing Kit (Oxford Nanopore, England), according to manufacturer’s protocols, and sequenced in a MinION Mk1C sequencing platform (R10.4 flowcell, Oxford Nanopore, England) at the Bioproduction Laboratory (INIA), Uruguay. We used Guppy v.5.0.11 in the “high accuracy” mode to base-call Oxford Nanopore Technology (ONT) reads and removed all reads under 1Kb for further analysis. We also filtered reads for quality using default settings in MinKNOW.

### RNA-seq data in axenic culture and during infection of *Piezodorus guildinii*

*B. bassiana* strain ILB308 was grown in different culture conditions for RNA extraction. To obtain axenic medium samples, ILB308 was grown on minimal medium (MM) during 4 days at 25°C after which fungal mycelium was collected from the plates and stored at −80 °C until RNA extraction.

To obtain samples of ILB308 during infection, it was previously grown in MM or MM supplemented with HC15 (MM+HC15). Fungal spores were collected from these two culture media plates and spore suspensions were prepared for insect inoculation according to Sessa et al. (2022). *P. guildinii* uncopulated females were inoculated by immersion in a conidial suspension of 1×10^7^ conidia/ml and maintained on an incubation chamber with 14/8hrs light/dark, at 25°C with 70% humidity. At 4 days post inoculation (dpi) living *P. guildinii* individuals were removed from the incubation chamber and stored at −80 °C until RNA extraction. RNA extraction was performed using the tissue, and the abdomen and thorax contents of dissected insects. Each sample (axenic cultures and inoculated insects) was ground with a RNAse-free mortar and pestle pre-cooled with liquid nitrogen, and total RNA was extracted using RNAeasy Plant isolation kit (Qiagen, Germany). Polyadenylated mRNA was strained from the total RNA and cDNA libraries were prepared using the Truseq RNA Library Prep mRNA kit (Illumina, San Diego, USA) following the manufacturer’s instructions. Constructed libraries were sequenced at the 150 bp PE with Illumina Novaseq6000, performed at Macrogen Inc, Korea.

### Genome assembly and gene prediction for *Beauveria bassiana*

Due to variations in genome coverage and potential structural disparities between *B. bassiana* strains, six different assembly approaches were employed to attain optimal assemblies for each strain. Sequences acquired from the Illumina platform were trimmed using Trimmomatic v0.36 [74] to eliminate low-quality and adapter sequences, and trimmed reads were assembled with SPAdes 3.5.0 [75]. Subsequently, short reads with basecalled ONT data were assembled with “SPAdes hybrid assembly” approach. The selection of k-mer lengths for assembly was automated based on read length, and contigs shorter than 200 bp were excluded. A third approach consisted of assembling basecalled ONT data using Canu v2.2 [76] with default parameters, employing the “genomeSize=36m” option based on the expected genome size [77]. Another hybrid assembling method was implemented using MaSuRCA v4.1.0 [78] with default parameters, merging Illumina trimmed reads with basecalled ONT data as the fourth approach and a consensus assembly was derived from the four previous assemblies using Flye v 2.9.2 [79] with specific parameters: “–-subassemblies -g 36m –-scaffold-no-alt-contigs” as the fifth. Lastly, an enhanced consensus assembly was generated by combining the previous Flye v2.9.4 [79] consensus assembly with POLCA polishing [80], followed by Samba scaffolder [81].

The quality of each assembly was assessed using the Quality Assessment Tool (QUAST 4.5) [82], with parameters –fungus, --split-scaffolds, and -min-contig=1000. Additionally, assembly completeness was evaluated using Benchmarking Universal Single Copy Orthologs (BUSCO) v5.0.0 [83] with – genome mode and the ascomycota_odb10 database. Performance of the different assembly approaches a_k_ ∈ {a_1_, …, a_6_} on the three data sets d_i_ ∈ {d_1_, d_2_, d_3_} using six pre-selected metrics m_j_∈{m1, …, m6} was performed and normalized according to Hölzer and Marz, 2019 to establish a matrix score (MS). These six metrics were number of contigs, size of largest contig, N50, % of mapped back reads, N’s per 100kb, and % of Complete BUSCO genes. For each combination of dataset d_i_ and metric m_j_, we defined a vector v^i,j^ of raw scores 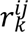 for each assembly tool a_k_ as: 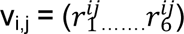. Then, we normalized the values of the vector v_i,j_ to the interval (0,1) using: normalize 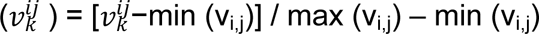,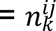. We defined the MS for an assembly a_k_ and a data set d_i_ as the sum of all (0,1)-normalized scores 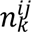 over the six pre-selected metrics m_j_ over a maximum MS of six.

The selected genome assembly for each strain underwent further manual curation by incorporation of telomeric regions with Teloclip v0.0.3 (https://github.com/Adamtaranto/teloclip); telomeric regions were added if supported by >10 raw ONT sequences. To assess the number of telomeric sequences and eliminate low coverage scaffolds (<5x coverage), we used Tapestry v1.0.1 (https://github.com/johnomics/tapestry.).

The final genome assemblies were structurally annotated with Funannotate v1.8.9 (https://doi.org/10.5281/zenodo4054262). Axenic RNA-seq reads (filtered, see below for details) were used as transcript evidence for gene predictions and the predicted protein-coding genes of *B. bassiana* strain ARSEF8028 (NCBI GenBank Acc GCA_001682635.1) were used as protein evidence. RNA-seq data for *B. bassiana* ILB299 was obtained from NCBI BioProject PRJNA991123 (Sessa et al., 2024).

### Comparative genomics and phylogenetic relationship of *Beauveria bassiana* strains

Genome-wide comparison between *B. bassiana* strains ILB205, ILB299, ILB308, and ARSEF8028 was performed by aligning genome assemblies using minimap2 [85] with parameters “-ax asm20” and plotted using D-genies (Cabanettes and Klopp, 2018). Circular visualization of genomes were performed using TBtools-II [87]

Orthofinder2 (Emms and Kelly, 2019) was used to identify orthologous groups of protein-coding genes among strains using the longest protein sequence as a proxy per gene. Orthologous gene pairs were identified based on amino acid sequence similarity and applying a coverage cutoff of 50% of the total length of the shorter gene sequence. Sequence similarity was established with BLASTp [89] and a threshold E-value 1e^−^5. For phylogenetic analysis, orthologous groups of five NCBI publicly available genome annotations of *B. bassiana* strains ARSEF2860 (GCF_000280675.1), ARSEF8028 (GCA_001682635.1), JEF-007 (GCA_002871155.1), ERL836 (GCA_010099065.1), D1-5 (GCA_000770705.1), our own three strains (ILB205, ILB299, and ILB308), and the outgroup species *Beauveria brongniartii* RCEF 3172 (GCA_001636735.1) were included. Single-copy core genes were aligned using MAFFT (Katoh and Standley, 2013) with option -maxiterate 1000. Phylogenetic tree of each orthogroup was deduced by software RaxML-NG [91] in GTRGAMMA model and 1000 bootstrap with Maximum likelihood (ML) method [92]. The final rooted tree was inferred by STAG method implemented in Orthofinder2 (Emms and Kelly, 2019).

### Functional annotation of protein-coding genes

Predicted protein-coding genes in the three *B. bassiana* strains were functionally annotated using BLASTp [89] against the non-redundant (nr) database of the National Center for Biotechnology Information (NCBI) with an E-value cutoff of ≤1e-5. The InterProScan suite [93] (accessed June 2023) was run with default parameters for the functional analysis of proteins, and eggNOGG [94] (accessed June 2023) was used for Kyoto Encyclopedia of Genes and Genomes (KEGG) assignments. All results were combined using Blast2GO [95] for the final functional annotation and Gene Ontology (GO) terms assignment, both using default parameters.

Candidate virulence proteins (CVP) were annotated based on at least one of the following five attributes: a) predicted secreted proteins (SP) identified using Phobius web server (https://phobius.sbc.su.se/) and SignalP v5 [96]. To exclude membrane associated proteins, we used TMHMM 2.0 web server (https://services.healthtech.dtu.dk/services/TMHMM-2.0/) with maximum 1 PredHel within the first 60 amino acids in combination with transmembrane domain prediction by Phobius web server (no TM predicted); b) Small SP, i.e., smaller than 300 aa; c) Secreted effector-like proteins identified with EffectorP3 (Sperschneider and Dodds, 2022); d) Proteins showing homology based on BLASTp to known virulence factors from the PHI-base (http://www.phi-base.org/). Results with e-value < 1e-5 and query coverage >60% were retained and those with ‘unaffected virulence’ PHI classification were removed; e) Identified carbohydrate-active enzymes (CAZymes; DbCAN2 (http://cys.bios.niu.edu/dbCAN2/)) (e-value < 1e−10) using HMMER [98], Hotpep [99], and DIAMOND [100]. Finally, because proteins interacting with host cells are mostly secreted, we specifically searched for those secreted CVP (SCVP) proteins that have at least two annotations: secreted and another CVP prediction.

To detect genes that could be part of secondary metabolite biosynthetic clusters, we used Antismash v7 (https://fungismash.secondarymetabolites.org) using relaxed strictness with all features enabled.

Moreover, we created a custom database of known *B. bassiana* virulence genes based on literature information, which included genes that had been functionally verified as involved in virulence in *B. bassiana*.

### Fungal transformation and morphology visualization

The expression of enhanced green fluorescent protein (*eGFP*) in ILB308 was achieved through the integration of the binary vector pCGEN-GFP [101], which contains both the geneticin resistance gene (*neo*) and a *eGFP* [102], using the *Agrobacterium tumefaciens*-mediated transformation method [103]. The transformed ILB308 was selected by 700ug/ml geneticin resistance in CPK medium and examined for GFP signals under a LSM ZEISS 800 confocal microscope with a 63x NA 1.4 Plan Apo objective. For GFP detection, a 488 nm laser line was used for excitation and a spectral range of 510-530 nm for emission with a GaAsP detector to enhance sensitivity. The Master Gain, Gain, and offset were kept consistent for all images captured and set below the levels used for controls without GFP. To visualize the fungal morphology at each stage, a transmitted channel using a 640-laser line was employed.

Development assays for morphology control between wild type and GFP-labelled strain were conducted using a conidial cell suspension aliquot (1 μl 1×10^7^ conidia/ml) spotted onto PDA plates containing: NaCl (1 M), sorbitol (1 M); H2O2 (1 mM) and Calco Fluor White (CFW) (200 ng/μl). Plates were incubated for 7 days at 25°C and the diameters of the fungal colonies measured. Standard PDA plates inoculated in the same way were included as controls.

### Bioassays and visualization of ILB308 strain during infection of *Piezodorus guildinii*

Virulence assays towards *P. guildinii* were made according to Sessa et al (2022) with minor modifications. Briefly, ten laboratory-reared adults of *P. guildinii* insects (five males and five females) were inoculated with a 1×10^7^ spore suspension of ILB308 or GFP labelled ILB308. Insects were immersed in the spore suspension for 10 seconds and immediately placed individually on a 55mm Petri dish containing sterile paper, water, and soybean grains for insect nutrition during incubation. Controls were immersed in water/Tween-saline solution. All treated insects were incubated at 24 ± 1°C for 11 days and insect mortality was assessed and recorded daily. The experiment was conducted per triplicate and cumulative survival curves were constructed.

For visualization of the infection process, a total of 30 female individuals were inoculated with GFP-labelled ILB308 and five living individuals at each time point between 1-5 days post inoculation were obtained for subsequently analysis. Fluorescent-transillumination imaging was conducted to visualize the overall distribution of ILB308-GFP labelled within the body regions of *P. guildinii.* A FLA 9000 biomolecular imager equipped with a blue laser and detectors compatible with GFP spectra was used. An initial image with low PMT signal (below 300) was captured for all insects to determine their body outline. Subsequently, a new image of the same insects was taken with high PMT signal (over 500) to identify the GFP-saturated regions. Both images were then combined using Fiji-Just-Image J software v1.54h (https://imagej.net/ij/). The overall distribution of GFP-labelled ILB308 throughout the body regions of *P. guildinii* was calculated as the percentage of GFP-saturated signal within the body outline.

### RNA-seq expression analysis

All raw RNA-seq data were filtered using Trimmomatic v0.36 [74] by trimming Illumina adaptors and bases with a quality score value less than 25 from both read ends (final minimum size allowed after trimming was 75 bp). Filtered reads were aligned to final ILB308 genome assembly using STAR [104], and RSEM (Li and Dewey, 2014) was used to determine read counts per gene. All statistical analyses of the expression data were performed using EdgeR v3.28 [106], and differentially expressed genes were identified using the exact test with the following conditions: log2Fold change of |1| and FDR (P< 0.05). Moreover, functional annotation of GO and KEGGs was used for enrichment analysis of DEGs using Fisher Exact tests.

### Availability of data and materials

The genomic sequences and RNA-seq datasets generated during the current study are available at NCBI BioProject repository, under accession number <u>PRJNA1117794</u>. Structural and functional annotations obtained in this study are available from the corresponding author upon reasonable request.

## Supporting information

**S1 Table.** Comparative evaluation of seven genome assemblies of Beauveria bassiana ILB205 based on seven quality assembly metrics and their correspondent normalized scores.

**S2 Table.** Comparative evaluation of seven genome assemblies of Beauveria bassiana ILB299 based on seven quality assembly metrics and their correspondent normalized scores.

**S3 Table.** Comparative evaluation of seven genome assemblies of Beauveria bassiana ILB308 based on seven quality assembly metrics and their correspondent normalized scores

**S4 table.** Summary of FungiSmash biosynthetic clusters prediction in four *Beauveria bassiana* strains

**S5 Table.** Structural annotation statistics for each scaffold in the ILB308 genome

**S6 Table**. List and composition of orthogroup that comprise genes present in ILB308 putative supernumerary scaffolds

**S7 Table**. Mapping stats from eight RNA-Seq libraries derived from three conditions of ILB308 in axenic and during infection of *Piezodorus guildnii*

**S8 Table**. Summary of GO enrichment analysis based on a Fisher exact test in ILB 308 during infection at 4dpi (vs MM axenic)

**S9 Table**. Summary of KEGG enrichment analysis based on a Fisher exact test in ILB 308 during infection at 4dpi (vs MM axenic)

**S10 Table**. Identification of known Beauveria bassiana virulence factors and the respective homologues in ILB308 strain

**S11 Table**. List and description of genes that were up regulated during infection at 4dpi in ILB308 present in predicted supernumerary scaffolds

**S12 Table**. Identification, description and expression values of genes up regulated during infection at 4dpi after growth in media supplemented with n-pentadecane

**S13 Table**. Identification and description of genes related to oxidoreductase activity up regulated during infection at 4dpi after growth in media supplemented with n-pentadecane

**S1 Figure.** *Beauveria bassiana* strain ILB308 core scaffolds.

**S2 Figure.** Macro and micromorphology analysis show no differences between wild type and GFP labelled strain.

## Notes

### Competing Interest Statement

The authors have declared no competing interest.

